# Contextual connectivity: A framework for understanding the intrinsic dynamic architecture of large-scale functional brain networks

**DOI:** 10.1101/068320

**Authors:** Rastko Ciric, Jason S. Nomi, Lucina Q. Uddin, Ajay B. Satpute

## Abstract

Investigations of the human brain’s connectomic architecture have produced two alternative models: one describes the brain’s spatial structure in terms of localized networks, and the other describes the brain’s temporal structure in terms of whole-brain states. Here, we used tools from connectivity dynamics to develop a synthesis that bridges these models. Using task-free fMRI data, we investigated the assumptions undergirding current models of the connectome. Consistent with state-based models, our results suggest that localized networks are superordinate approximations of underlying dynamic states. Furthermore, each of these localized, moment-to-moment connectivity states is associated with global changes in the whole-brain functional connectome. By nesting localized connectivity states within their whole-brain contexts, we demonstrate the relative temporal independence of brain networks. Our assay for functional autonomy of coordinated neural systems is broadly applicable across populations, and our findings provide evidence of structure in temporal dynamics that complements the well-described spatial organization of the brain.

---

A major endeavor in neuroscience is to characterize the spatiotemporal organization of the brain into functional systems^1^. By identifying patterns of synchronous brain activity, functional magnetic resonance neuroimaging (fMRI) techniques have partitioned the human brain into large-scale networks^2,3^. These functional networks are stable across individuals and populations^4–6^, are roughly consistent across task-evoked and task-free states^7,8^, and are present across mammalian species^9,10^. A hierarchically modularized set of canonical networks is now widely accepted as an organizational principle of the brain^11,12^. Indeed, an expanding literature relates networks to specific psychological functions and individual differences^13–16^, with the potential for improved clinical diagnosis or treatment outcome metrics^17–19^.

However, recent work has called into question how accurately this canonical network model represents underlying neural architecture. In particular, many methods used to delineate networks rely on two implicit assumptions. First is the spatial assumption that each brain region participates in exactly one network. Casting doubt on this are recent models suggesting that brain regions can engage with several different networks^20–22^, dynamic causal models showing that connectivity between brain regions changes as a function of the experimental context^23^, and graph theoretic models intimating the existence of neural hubs that recruit multiple networks^24–27^. Second is the temporal assumption that the connectivity within each network remains relatively stable during fMRI experiments. This, too, has been called into question, with recent work^28^ suggesting that the brain is dynamically multistable. That is, the brain may occupy any of a number of connectivity states over time, each with a distinct network architecture^29^. Such a multistable model has been applied to discover novel biomarkers for pathology^30–32^ and to track changing cognitive demands^33^.

Currently, it is unclear whether these recent findings reflect modest fluctuations nested within the canonical network architecture, or whether the spatial and temporal assumptions of commonly used network models must be re-evaluated. In the present study, we evaluate the stability, homogeneity, and independence of six networks to determine whether each network’s temporal variability constitutes stochastic variation about its time-averaged structure, or whether it reflects temporally distinct network connectivity states (or NC-states). We identify putative NC-states using k-means clustering of connections computed over a sliding window^28^ (Figure 1). We next characterize the whole-brain milieu in which NC-states occur, identifying novel organizational principles from which to further understanding of the brain’s intrinsic architecture. Our findings diverge from assumptions of spatiotemporal stability, and suggest instead that canonical networks are superordinate representations of several NC-states, each of which describes the state of a network at a given moment in time and is associated with a distinct whole-brain context. Surprisingly, we also find that individual network connectivity is relatively independent of whole-cortex connectivity. Based on these findings, we develop a novel organizational framework, *contextual connectivity*, towards reconciling network- and state-based models of the human brain.

**Figure 1.**
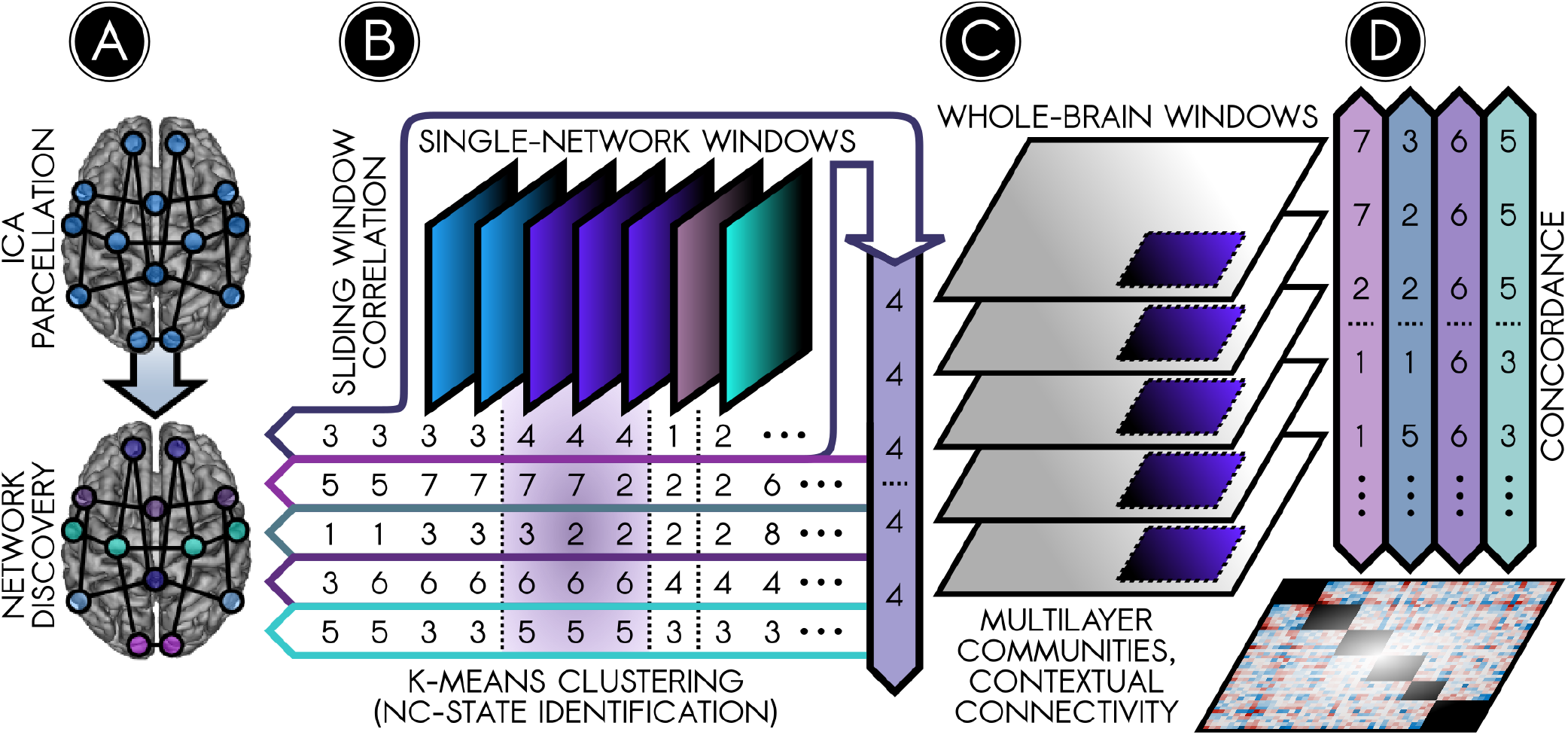
Summary of analysis steps. **(A)** ICA was used to parcellate the human brain into 70 cortical and 10 subcortical/cerebellar nodes (top). Cortical nodes were assigned to six canonical functional networks using a community detection algorithm (bottom; results in Figure 2). **(B)** Temporal fluctuations in connection strength between nodes were identified using sliding-window correlations. A k-means clustering analysis identified prominent, recurring ‘network connectivity states’ (NC-states) among nodes within each canonical network. The top row schematically illustrates four recurring NC-states of one canonical network. The bottom rows show the NC-states in other networks for corresponding time windows (results in Figure 3a). To see how NC-states in a given network relate to the connectivity state of the rest of the brain, two approaches were taken. One approach averaged the connectivity matrices over a given NC-state’s time windows (such as for NC-state 4 for the network in the top row in the schematic) to determine **(C)** the whole-brain connectivity context (WBCC, grey portion of matrix) in which each NC-state (purple portion) occurred (results in Figure 3b). The other approach examined synchrony between NC-states in different canonical networks using a Bayesian concordance matrix **(D**, *bottom*) to test whether NC-states in different networks *(top)* relate to one another over time (results in Supplementary Figure 4).

## Results

### Canonical network identification

The spatial segregation of cortical parcels into functional brain networks forms the basis of modern analyses of the human connectome. Here, we reproduced this canonical network partition as the first step toward a revised understanding of networks as temporally dynamic entities. Specifically, we used spatial independent component analysis (ICA)^34^ to localize 80 sources of variance in the BOLD signal. A connectome was defined using the 80 components as nodes and using the Pearson correlation coefficients between node timeseries as evidence of connection strength^35^. Nodes were assigned membership to canonical brain networks by training a community detection algorithm^36,37^ using an *a priori* hypothesis^3,38^. We thus obtained six communities of nodes exhibiting one-to-one correspondence with six reference brain networks: the visual network (VIS), the somatomotor network (SOM), the dorsal attention network (DAT), the cingulo-opercular/salience network (SAL), the executive control network (EXE), and the default mode network (DMN)^2,3^. Spatial cross-correlations between these communities and reference networks ranged from approximately 0.49 to 0.68, well above a previously established cutoff^38^, a positive indication that the canonical partition was reproduced (Supplementary Figure 1C). Figure 2a illustrates the clear spatial correspondence between the networks of our partition and canonical functional networks^2,3,28^. Figure 2b depicts the time-averaged connectome; the majority of strong and specific correlations among nodes were localized to within-network connections. We used this partition to organize subsequent analyses examining the moment-to-moment connectivity of functional networks.

**Figure 2.**
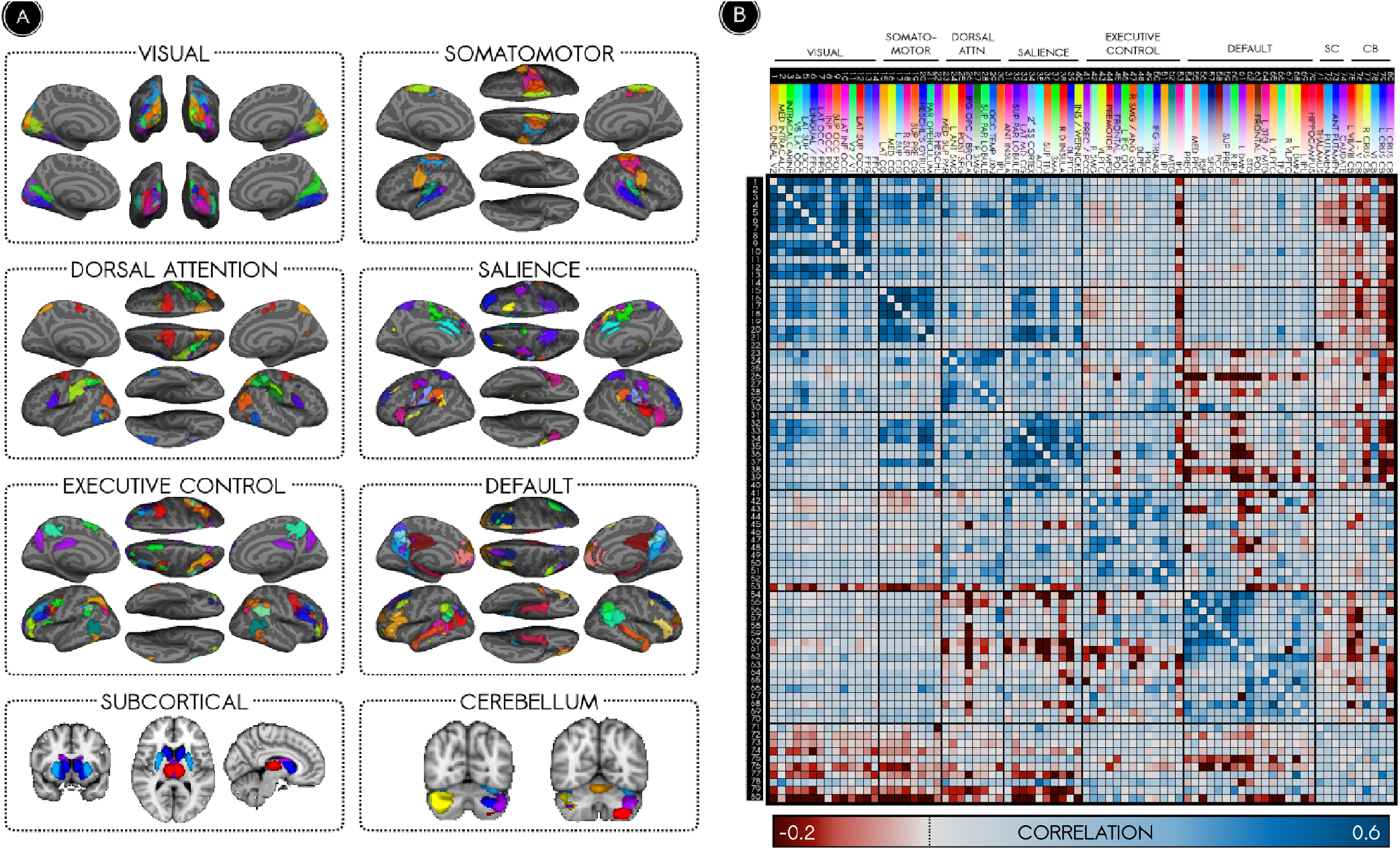
Canonical functional networks of the time-averaged connectome. **(A)** Spatial maps of nodes are illustrated according to their membership in functional networks. **(B)**Strengths of connections between nodes are illustrated in a whole-brain adjacency matrix. The strongest connections of the time-averaged functional connectome occur among nodes of the same network.

### Each canonical network is resolvable into a set of network connectivity states (NC-states)

While the brain’s organization into networks ranks among the most robust findings in functional connectivity, the stability of brain networks over time is under dispute. Recent evidence suggests that the moment-to-moment synchrony among the regions constituting each functional network may deviate markedly from that network’s canonical structure^28^. Here, we evaluated this hypothesis. For each of the six networks, we calculated the extent to which different ‘network connectivity states’ (NC-states) could arise among nodes in the network. We began by using k-means clustering to identify putative NC-states for each canonical network. Each NC-state repsented a distinct pattern of synchrony among the nodes within a network. Compared with expectation under permuted and phase-shifted null models, the NC-states we observed were well-differentiated (*p*<0.01) and represented distinct clusters (*p*<0.01; Supplementary Figure 2). Figure 3a illustrates the NC-states of the DMN sorted in ascending order of intrinsic connectivity.

**Figure 3.**
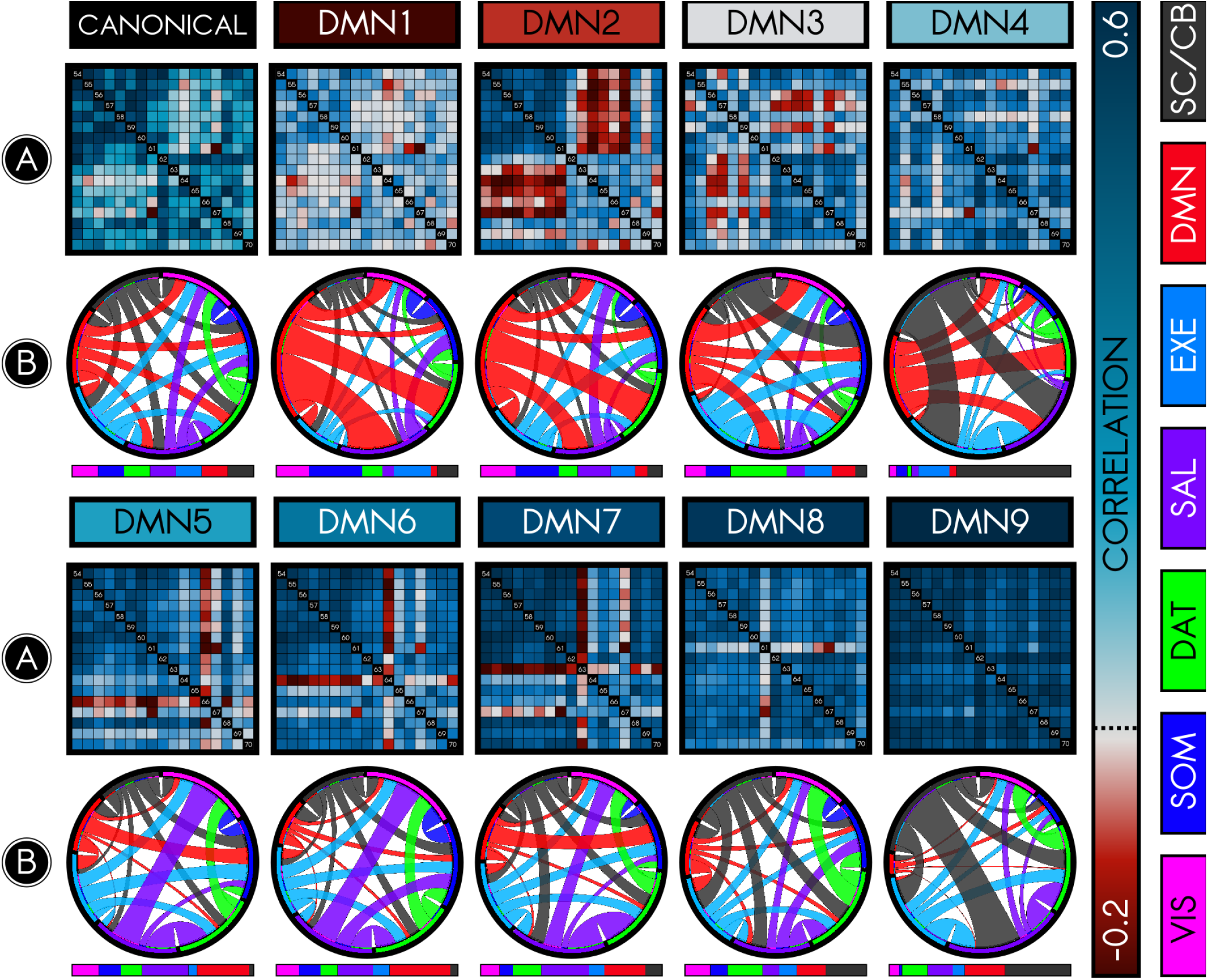
Connectivity dynamics of the default mode network (DMN). Dynamic functional connectivity analysis among nodes of the DMN revealed that the DMN is composed of several network connectivity states (NC-states). This figure summarizes the connectivity patterns of the canonical (time-averaged) DMN (*first row, left*) in comparison with the 9 observed NC-states (DMN1-9), organized in order of low intrinsic connectivity (dark red) to high intrinsic connectivity (dark blue). **(A)** illustrates the connectivity (Pearson correlation) among the 17 nodes in the DMN, either time-averaged or for specific NC-states. Each NC-state exhibits a distinct profile of connectivity among nodes. **(B)** illustrates relative within- and between-network allegiance for each NC-state and its whole-brain context. Allegiance is the probability that nodes are in the same community when that NC-state is present. The circle plot illustrates *between-network* allegiances for each NC-state relative to the time-averaged connectome. The color codes along the rim signify different canonical networks as labeled at right (e.g., red for DMN). Longer rim segments indicate that the network has greater allegiance to other networks relative to the time-averaged state, and the size of connections between rim segments indicate strength of between-network allegiances. The time-averaged circle plot defines a ‘baseline’; thus, each rim segment is of equal size. The remaining plots show that allegiance between networks changes across NC-states (to emphasize changes in allegiance, plotted allegiance ratios were rescaled exponentially). The adjacent bar illustrates *within-network* allegiance using corresponding color codes. Longer segments indicate that the network has greater intrinsic allegiance, again relative to the time-averaged ‘baseline’. For instance, DMN in DMN1 exhibits weak within-network allegiance but strong between-network allegiance, which appears to be driven by greater allegiance to DAT, SAL, and VIS. DMN’s intrinsic allegiance increases from DMN1 to DMN9. Conversely, DMN’s allegiance to other networks substantially drops from DMN1 to DMN9.

Inconsistent with models that suggest uniformity of brain networks over time, we found that networks could be resolved temporally into NC-states. Remarkably, no NC-state was completely predominant for any network. Instead, all subjects exhibited multiple NC-states per scan, suggesting that the many NC-states observed per network were not idiosyncratic to particular subjects. For the default mode network, for instance, between 7 and 9 NC-states were represented in the majority of subjects, with no subject exhibiting fewer than 4. Moreover, gross features of NC-states were conserved at the single-subject level, and the majority of NC-states replicated across split-half samples (.88 feature-wise correlation, 36 out of 46 NC-states replicated; Supplementary Figure 3A). Taken together, these results indicate that each brain network is incompletely characterized by its time-averaged connectivity profile.

### NC-states occur in specific whole-brain connectivity contexts (WBCCs)

We used NC-states as the building blocks of a framework for evaluating the *temporal independence* of canonical networks. We first posed the question: Given information about the state of one brain network, can we make inferences about the state of another network, or of the brain as a whole? To answer this question, we fashioned hypotheses based on whole-brain connectivity contexts (WBCCs)^39^. The WBCC of each NC-state was defined as the average (whole-brain) environment in which that (single-network) NC-state was present. We used multilayer community detection^37^ to obtain an allegiance matrix^40^ representing each NC-state’s WBCC. In an allegiance matrix, the weight of the edge connecting a pair of nodes represents the probability that those nodes will be found in the same community over all time windows during which that NC-state is present. Allegiance matrices, unlike correlation matrices, are more sensitive to specificity than to magnitude of connections. We first tested a model of *complete temporal independence*: the null hypothesis that an NC-state’s WBCC did not significantly differ from the time-averaged connectome, which would suggest that each network’s connectivity state changed independently of the remainder of the brain. In contrast with the null model derived under this hypothesis, all NC-states were associated with specific changes in the functional architecture of the whole brain (Supplementary Figure 3B). Figure 3b uses a Circos plot to illustrate changes in the WBCC associated with each NC-state. For example, DMN4 was associated with increased allegiance among salience, executive, and subcortical systems, while DMN7 was associated with increased allegiance between the salience and dorsal attention networks. In supplementary analyses, we also tested interdependence between individual networks in another way by using a Bayesian *concordance* metric. This showed that the occurrence of a state in one network was predictive of the occurrence of particular states in other networks (Bonferroni-adjusted *p* < 0.05, Wilcoxon test; Supplementary Figure 4B). These results are inconsistent with temporal independence of brain networks, and instead support the view that information about the state of the entire brain is embedded in each network.

### Dynamically determined NC-states recapitulate time-averaged subnetworks

The analyses above suggest that brain networks are temporally resolvable into transient NC-states. However, it is not yet clear that this complexity adds notable value. To address this, we first examined whether the NC-states we observed using dynamic analyses recapitulate prior results from static connectivity analyses. Such work has shown that the DMN is composed of two subnetworks, one anchored in the medial temporal lobe (MTL) and the other in the dorsomedial prefrontal cortex (DMPFC), both of which converge onto a ’midline core’ (Figure 4A, inset). We first examined whether our analysis reproduced these subnetworks of the DMN. We used hierarchical clustering to determine whether a network’s nodes associated into subnetworks (Supplementary Figure 5). NC-states DMN2 and DMN3 showed a clear correspondence with previously reported MTL and DMPFC subnetworks, respectively. DMN2 featured enhanced connections among medial nodes (red and violet), while DMN3 featured enhanced connections among lateral nodes (blue and violet). The precuneus, PCC, medial PFC, and right IPL – nodes that remained cohesive in both NC-states (violet in Figure 4a) – map onto the midline core of the DMN following prior work^41^ (Figure 4A, inset). These findings validate our dynamic approach insofar as it captures known findings from static connectivity studies.

**Figure 4.**
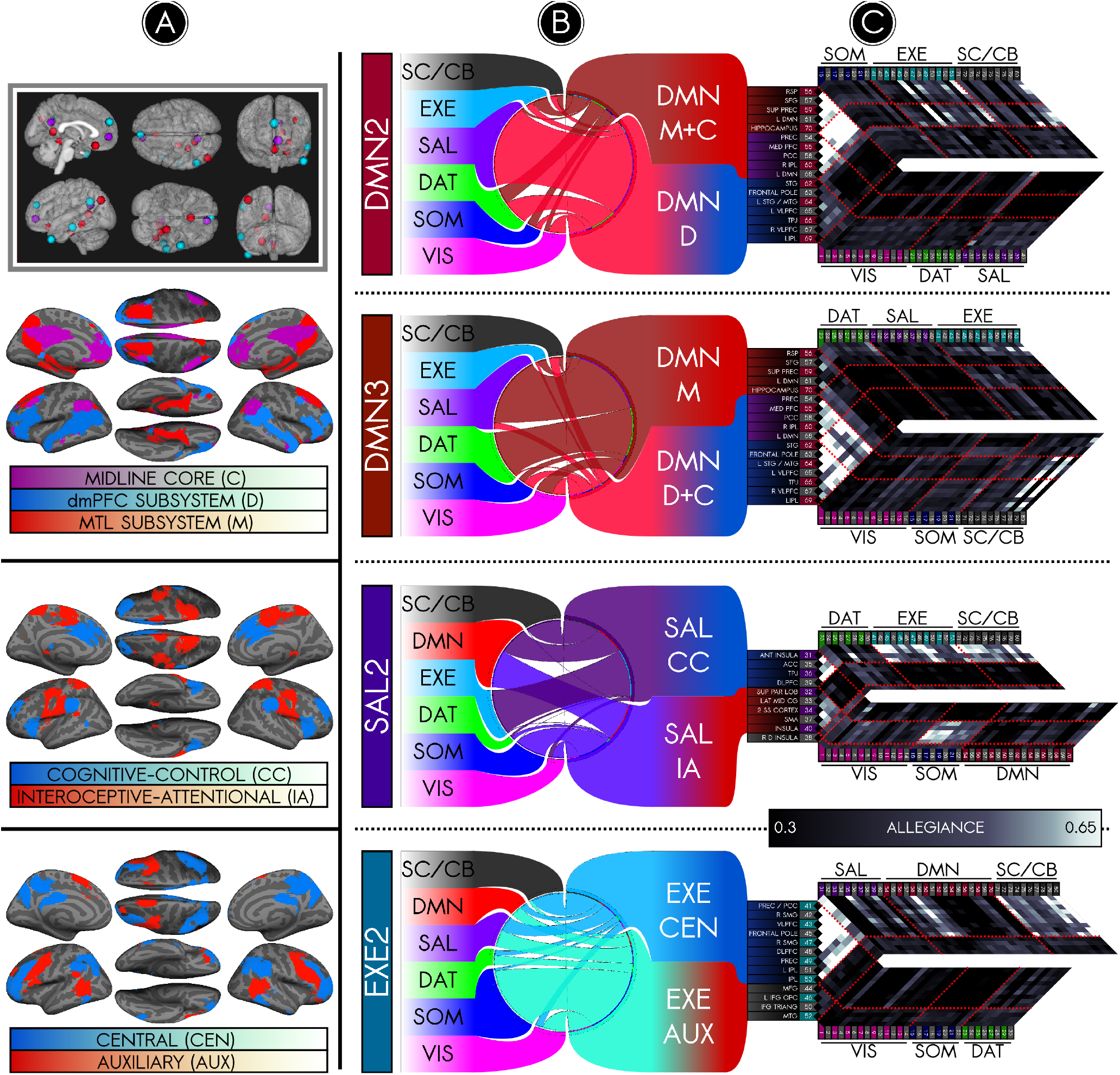
The salience, executive, and default mode networks dynamically segregate into temporally decomposable but spatially overlapping subsystems. **(A)** *Top to bottom*: Spatial maps of dynamically segregated subsystems of SAL, EXE, and DMN. **(B)** Circle plots illustrate changes in subsystem connections relative to the time-averaged connectome. **(C)** Allegiance matrices illustrate that nodes within different subnetworks exhibit differential allegiance to external networks. *Top*: The default mode network exhibited two NC-states that reflected its underlying connectomic architecture. DMN2 featured selective cohesion in an MTL subsystem (M), while DMN3 was characterized by selective cohesion in a dmPFC subsystem (D). Several medial core regions (C) remained cohesive in both states. *Inset, top left*: Dynamic DMN subsystems recapitulate subsystems of DMN previously reported using a time-averaged connectivity analysis^15^. *Third row*: NC-state SAL2 was characterized by a bifurcation of the salience network into a cognitive-control core (CC) and an interoceptive periphery (IA), each exhibiting distinct and antagonistic connectivity profiles. CC preferentially linked into control systems, while IA preferentially linked into perceptual systems. *Bottom*: NC-state EXE2 was characterized by enhanced modularization of a central executive system (CEN) and demodularization of auxiliary nodes (AUX).

We then addressed two additional questions afforded by our dynamic approach. First, what is the status of the decoherent DMPFC nodes during DMN2, and of the MTL nodes during DMN3? Our findings indicate that in both cases the decoherent nodes demodularize (Figure 4b-c). That is, the connectivity of the DMPFC subnetwork with other DMN nodes diminishes in DMN2, while its connectivity with other networks (e.g. SAL) increases. Similarly, the connectivity of the MTL subnetwork with other DMN nodes decreases in DMN3 while increasing with other networks (e.g. DAT and SAL). Second, we explored whether networks other than DMN also have NC-states that suggest the configuration of nodes into subnetworks. We observed NC-states corresponding to fractionation of the SAL and EXE networks (and replicated these in split-half samples). In SAL2, salience nodes assorted into two antagonistic subsystems (Figure 4a). We suggest putative functions based on their distinct connectivity profiles, but caution that a definitive functional description would require formal probing using tasks. SAL2 exhibited a ’cognitive-control’ core (blue) that became negatively correlated with an ’interoceptive-attentional’ periphery (red). While the ventral core remained selectively connected to EXE, the dorsal periphery connected strongly into SOM and VIS (Figure 4b-c). As for the EXE network, EXE2 was characterized by a splitting of the EXE into two sets of nodes (Figure 4b-c). While one set of nodes appeared to lose cohesion with the rest of the EXE and each other (red), another set of nodes exhibited increasing modularization (blue). The DMN subsystems coherent in DMN2 and DMN3 were similarly modularized (Figure 4b). These modularizations may reflect a shift toward specific, localized computation in these subsystems.

### Canonical networks are relatively independent

The preceding analyses provide evidence of interdependence among brain networks but do not test whether networks nonetheless retain a relative degree of independence. Indeed, it is notable that temporal independence of networks has largely been assumed on the basis of community structure rather than formally tested. We reasoned that for a given sample of nodes, the extent to which these nodes’ connections cannot explain dynamic connectivity among the remaining nodes is the extent to which these nodes are *independent*. Owing to the putative modularization of canonical brain networks, we hypothesized that dynamics of known networks would poorly explain the entire cortex’s dynamics in comparison with dynamics of a random sample of nodes, or a *pseudo-network*. We calculated how well the NC-states of canonical and pseudo-networks explained the temporal variation in the cortical connectome using the within-cluster sum of squares (WCSS) error metric. The WCSS error is a measure of the quality of a putative clustering solution, or the extent to which it explains the variability within a dataset; a higher WCSS error corresponds to a poor-quality clustering solution, which may be interpreted as evidence that a (pseudo-)network’s local connections poorly predict global connectivity; i.e., the (pseudo-)network is more independent. We proposed whole-cortex clustering solutions on the basis of only information from connections within each (pseudo-)network and calculated the WCSS error for each proposed solution. Consistent with our hypothesis, dynamics of canonical networks explained significantly less whole-cortex variation than did dynamics of pseudo-networks (Figure 5a; *p* < 0.05, Wilcoxon test). Furthermore, canonical network NC-states were significantly less concordant with cortical states than were pseudo-network NC-states (Figure 5b; *p* < 0.01 except SOM, Wilcoxon test). These findings suggest that (while fluctuations occurring on a dynamic level across networks are of importance) canonical networks nevertheless retain a degree of independence from the rest of the cortex.

**Figure 5.**
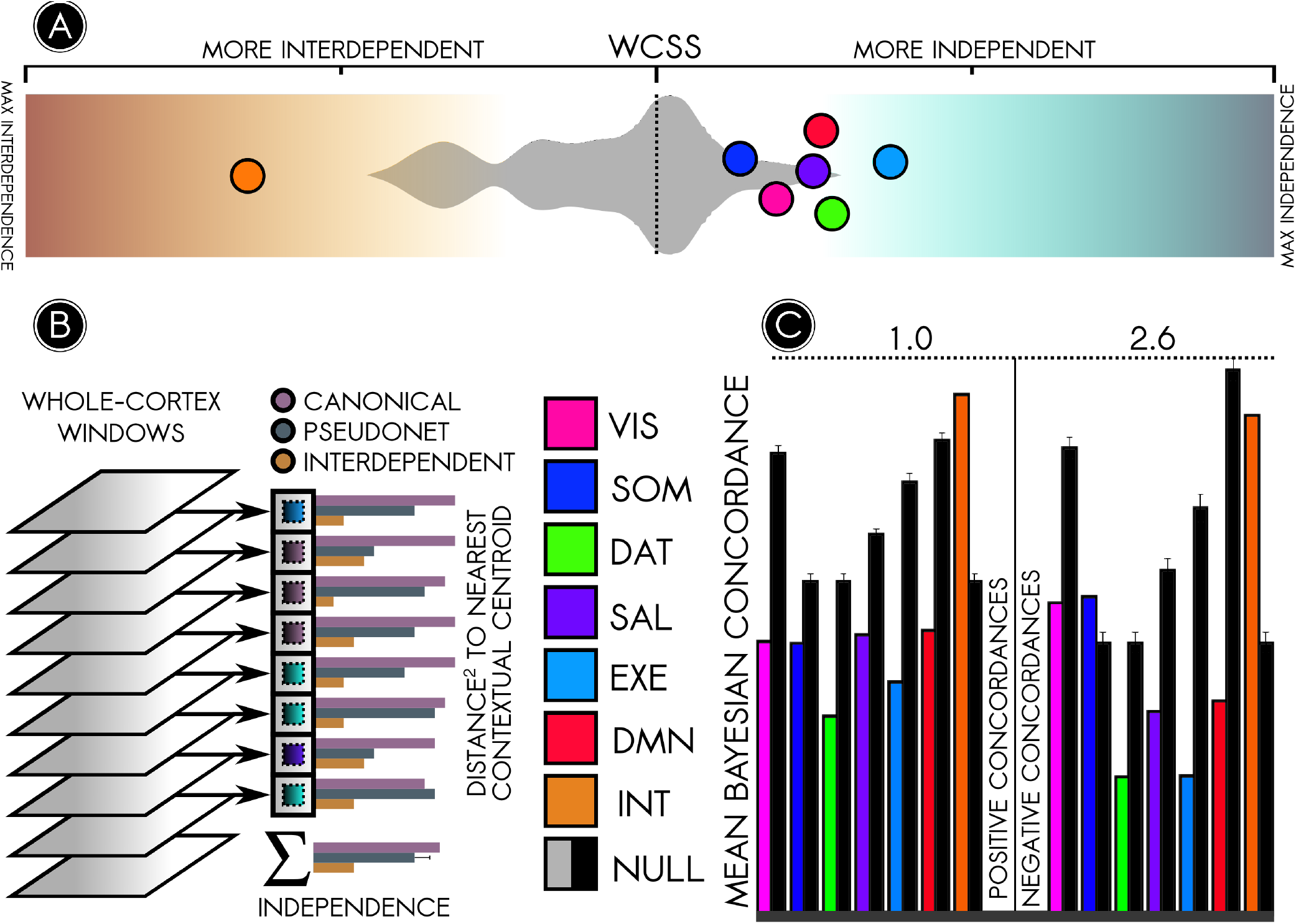
Canonical networks are significantly more temporally independent than random cortical subsystems. **(A)** The relationship between a canonical network’s intrinsic variation and whole-cortical variation can be quantified using the within-cluster sum-of-square errors (WCSS). The WCSS approach demonstrated that canonical networks (colored circles) were more independent than random pseudo-networks (grey silhouette, background) (*p* < 0.05, Wilcoxon test). **(B)** Schematic of the WCSS approach used to determine the independence of a canonical brain network. The whole-cortex connectivity pattern captured in each time window was matched to a context (contextual centroid) of a NC-state of the network under investigation. Matching was based on proximity in the brain’s connectomic state space. The squares of the distances separating windowed connectivity patterns from their matched contextual centroid were added together to determine the WCSS error; a higher WCSS error corresponded to greater independence. A null distribution was built by applying the same approach to random pseudo-networks. **(C)** A separate metric, the mean Bayesian concordance, recapitulated the results obtained using the WCSS error. Here, greater concordance corresponds to more interdependence. Each canonical network (colored bars) was compared against a null distribution of pseudo-networks (black bars); with one exception, canonical brain networks were significantly more independent (*p* < 0.01).

As an exploratory aim, we used these measurements to characterize a maximally interdependent brain subsystem. We hypothesized that such a ‘hub’ system would better explain cortical dynamics than would pseudo-networks and that it would exhibit NC-states highly concordant with cortical states. Several subsystems satisfied these criteria; of these, the most potent was the ‘most interdependent’ (INT) subsystem presented in Figure 6 (*p* < 0.01 for WCSS and concordance metrics). INT nodes included medial and lateral prefrontal cortices, midcingulate gyrus, middle temporal gyrus, fusiform gyrus, dorsal somatomotor cortex, and lateral occipital cortex. Unlike prior attempts to identify hubs, INT nodes were not characterized by high degree^18^ or participation coefficient^24^. Instead, they appear to be representative nodes of their parent networks and may lack substantial anatomical connections with one another. It is possible that traditional graph-based hubs entrain synchrony among INT nodes.

**Figure 6.**
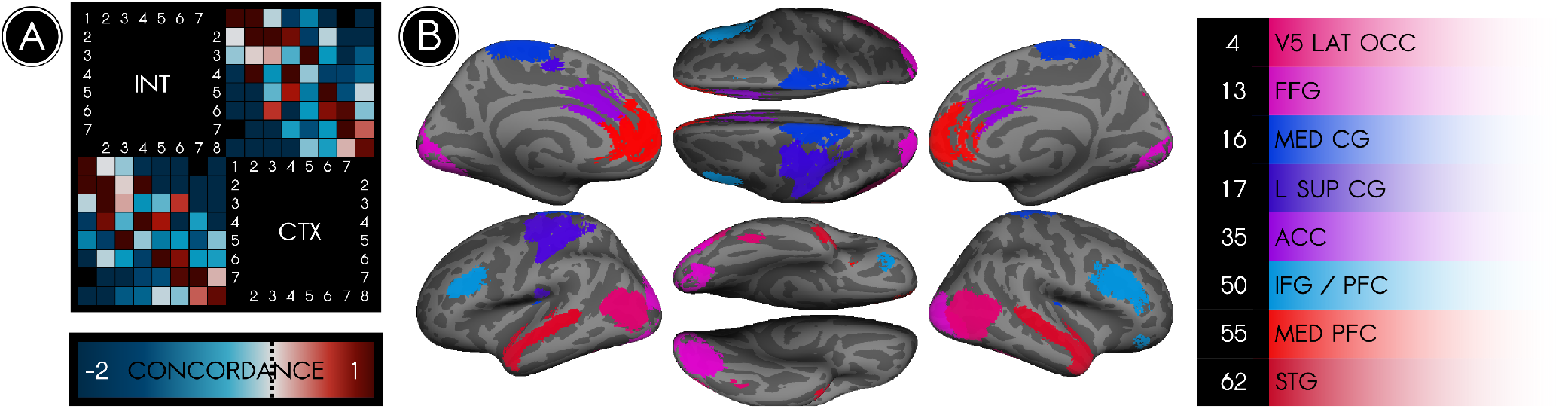
The most interdependent brain system consists primarily of connections between nodes in different networks. We identified an interdependent (INT) node set characterized by its high temporal interdependence with the cortex as a whole (CTX). **(A)** The Bayesian concordance between states of INT and states of CTX is represented as a state-by-state matrix; a positive concordance indicates that states co-occur more often than predicted by chance. Every state of INT was highly concordant (*concordance* > 1) with at least one state of CTX, reflective of similarities between the trajectories of INT and CTX through their respective state spaces. **(B)** The 8 nodes in the INT set, color coded according to their parent canonical network and listed at right. These nodes differed from graph-theoretical hubs; rather than being situated in areas where multiple networks overlapped, they were typically representative nodes of their parent networks. Because these nodes represented a diversity of brain networks, nearly all connections of INT were *between-network* rather than *within-network* connections.

## Discussion

An ideal model of the brain distills its dynamic, high-dimensional information^23^ into interpretable constructs without sacrificing fidelity. Towards this goal, research using functional connectivity has centered on two models. The first of these models emphasizes the spatial dimension, and parcellates brain activity into spatially localized functional networks^2,3,5^. A second model emphasizes the temporal dimension, and parcellates brain activity into dynamically recurring states^28,31,32,42^. While researchers have nominally acknowledged that the simplifying assumptions of each model introduce numerous deficiencies and limitations^20,28^ the preponderance of studies in functional connectivity continue to rely upon them.

In the present study, we deconstructed the elementary units of both models – the spatial network and the temporal state – into a ’common factor’, or a spatially localized connectivity state (i.e., a NC-state). We then evaluated the assumptions underlying network-based models, in particular the assumptions that brain networks are independent and stable. Inconsistent with the canonical model of localized and temporally stable networks, we found that brain networks are temporally decomposable into an array of possible connectivity states. Moreover, each local connectivity state also provides information about the global state of the entire brain. However, inconsistent with whole-brain models that disregard network boundaries, our results also indicate that networks retain a degree of modularization; connectivity patterns among nodes of a single network provide less information about the whole-brain dynamic state than do connectivity patterns among randomly selected nodes.

To accommodate these findings, we advance an alternative model, *contextual connectivity*, and a corresponding analytical framework. In our model, networks are better thought of as composed from a set of dynamically recurring states, each of which is associated with a specific whole-brain context. The combination of the network state and the whole-brain state (outside the network) provides a tractable balance that bridges two analytical levels of cognitive neuroscience: the spatially localized, temporally general network and the temporally localized, spatially general brain state. Thus, our model takes advantage of the spatial simplification provided by canonical networks, which is empirically supported by our findings, while also capturing dynamic reconfigurations of nodes as suggested by dynamic models.

(To be sure, this is not the only possible interpretation consistent with our evidence. An alternative model might be developed by first decomposing the whole brain dynamically and afterward identifying spatially localized, synchronous systems that recur over time. Thus, instead of identifying localized connectivity states, this approach would identify transient coalitions of nodes that manifest during particular time windows depending on moment-to-moment affiliations and disaffiliations among nodes. Such coalitions are likely to be, on average, roughly coterminous with network boundaries. While this alternative is consistent with our findings, it is unclear how to identify recurrent coalitions at this time.)

Analytically, our framework comprises four main steps, which in principle are extensible to any set of complex brain systems: (i) identify a system of interest (e.g., a canonical brain network), (ii) determine the internal temporal states of that system (here, using dynamic functional connectivity), (iii) determine the external contexts of those internal states (here, either by computing the average connectivity pattern of the whole brain or by identifying co-occurrent states of other systems), (iv) quantify the affiliation between the system’s internal states and external contexts (here, using the WCSS error or the concordance). Notably, our analytical framework provides a better opportunity for capturing multivariate or nonlinear exchanges between networks. This is because our model captures two levels of synchrony dynamics: not only node-to-node connectivity measured within a given temporal window, but also state-to-state concordance. At a particular time, it is possible that network A’s nodes have low moment-to-moment connectivity with nodes in network B, while the states of the two networks at the same time are highly concordant. Such cases indicate a complex interplay between brain networks that may be reflective of nonlinear or multidimensional interactions mong brain regions.

Because fMRI, and dynamic connectivity in particular, is susceptible to a number of spurious phenomena, we took caution to ensure that our results were driven by effects of interest rather than noise. Two artefactual processes are of particular concern in dynamic analyses of task-free fMRI: subject motion and sampling variability. First, in-scanner subject motion can bias connectivity results in favour of connections between regions that are close together in physical space^43,44^. We repeated our experiment on a low-noise subsample of the cohort, which we obtained by censoring epochs of high motion^44^. Overwhelmingly, we observed states nearly identical to those observed in the cohort as a whole, indicating that the dynamic effects we report were not explained by motion. Second, a recent body of work^45^ suggests that transient patterns of brain connectivity are not structured manifestations of a set of underlying brain states; instead, they are artifacts arising due to variable sampling of a single underlying state that is stable across time. To account for this possibility, we generated surrogate data by applying a randomized phase shift to the connectivity data, thus preserving the structure of such a stable underlying state but disrupting any organized dynamic connectivity patterns^45^. On the whole, we observed that connectivity states in the empirical data were better differentiated than those in the surrogate data, indicating that the connectivity patterns we observed reflected multiple underlying brain states. In addition to the analytical precautions taken, our replication of known neural architecture attests to the fidelity of our approach. First, we observed local connectivity states of the default mode network that mirrored previously reported task-related subsystems^15^. Second, our analysis of the independence of local connectivity states recapitulates the known spatial organization of the brain into functional networks^3^. The convergence of dynamic connectivity with previous results from the task-evoked and resting-state literature corroborates the argument that dynamic connectivity analyses are capable of detecting true neural architecture and not only spurious fluctuations.

### Implications for structure-function mapping

The identification of large-scale brain networks is motivated by the possibility of an improved mapping of brain structure to psychological function. However, the conjectured functions of large-scale brain networks are highly generalized and evade psychological intuition^13^; instead, they reflect the intrinsic degeneracy and pluripotency of the brain^46^. This lack of specificity suggests that brain networks are not the atomic ingredients of neural function. Indeed, analyses of static functional connectivity have revealed that large-scale brain networks are spatially dissociable into subnetworks^41^. Specific subnetworks can be selectively engaged using targeted task conditions^41,47^ suggesting that they support specific operations of their parent networks’ function. Here, we offer an update to this interpretation using dynamic, time-resolved approaches.

Patterns of neural activity identified as default mode subnetworks are recruited under specific task constraints^41,47^. We find that the same patterns also occur spontaneously and are detectable in dynamically occurring brain states (Figure 4). Moreover, our analysis provides additional insights and analytical metrics pertaining to subnetworks from a dynamic framework. First, we provide insight into how nodes – such as those comprising the MTL and DMPFC subsystems of the DMN – behave when one subsystem temporarily dominates over the cohesiveness of the network. In both cases, the decoherent subsystem appears to demodularize, increasing its connectivity to other networks. Second, whereas time-averaged approaches resolve networks into spatially non-overlapping subnetworks, our analysis permits spatial overlaps between nodes. As such, our approach is more capable of handling cases of degeneracy and pluripotency, which are considered to an important feature of complex systems^48^. Third, our approach allows us to probe the global context of each subnetwork as reflected in its NC-state (i.e. DMN2 or DMN3).

### Additional implications for functional connectomics

According to dynamical systems models, brain activity can be understood as tracing a trajectory through a multidimensional ’state space’. A question of current interest is how to identify the contributions of different brain regions to the brain’s overall trajectory. Our findings suggest that different types of regions contribute in distinct ways according to their network properties. Specifically, we found that INT nodes (Figure 6) provide maximal information about the status of the brain as a whole. In that sense, these nodes approximately denote the general location of the brain in its multidimensional state space. Notably, the connections of this system are representative linkages *between networks*. In contrast, the relatively independent *within-network* connections may provide more localized information about the brain’s status, or specific coordinates within its general location (Figure 5). These findings suggest that the state of the brain is coarsely determined by between-network connections, while within-network connections guide the brain to more specific states.

We also observed that the executive control network was consistently the most independent brain system across validation samples. Prior work has demonstrated that the executive network exhibits a nonspecific or global pattern of connectivity in the resting state^2,49^. This nonspecific pattern represents a temporal average over a highly variable dynamic repertoire of connections to all other networks^27^. The independence of the executive control network during the resting state indicates that its intrinsic activity is relatively unconstrained by activity across the remainder of the cortex. This property of the executive network may enable it to flexibly update its connections and steer the brain into a multitude of difficult-to-access states in response to changing cognitive loads^27,50^.

### Implications for individual differences

The discovery of large-scale functional networks has prompted considerable efforts to understand how these networks relate to individual differences. Prior work has focused primarily on whether canonical networks show topological variation across individuals. However, examining individual differences in network architectures requires a precise characterization of intrinsic connectivity networks. Here our study contributes in several ways. First, our findings suggest that canonical network analyses of individual differences run the risk of conflating subject differences in state topology with differences in network topology. Because each network can be resolved into distinct NC-states, it may be more informative to isolate states whose between-subject connectivity differences most strongly relate to individual difference variables. Second, our findings introduce novel approaches for relating differences between individuals to differences in network architectures. Dwell times^31^ and transition frequencies^51^ of whole-brain states have been identified as correlates of schizophrenia; analogous metrics computed for localized states could elucidate network drivers of pathology. Furthermore, the contextual independence metrics that we introduce might illuminate previously overlooked correlates of individual difference variables; specific pathologies may be reflected in a failure of systems to coordinate or, reciprocally, a failure of systems to segregate (elevated independence or interdependence). The methods we present for examining the degree of independence of brain systems could illuminate new relationships.

## Conclusions

Researchers have long acknowledged that brain networks are not immutable, monolithic entities, but analytical strategies consistent with this acknowledgment have been limited. Methods aimed at recapitulating canonical networks fail to capture important dynamics occurring within and between those networks. However, analyses that do away with network assumptions often present challenges of interpretability and complexity. Our approach, *contextual connectivity*, addresses this issue by introducing an intermediate level of analysis that not only respects the robust finding that networks are relatively autonomous, but also recognizes that networks are at best superordinate approximations of dynamically recurring states. As such, NC-states provide a tractable approximation of the functional connectome that maintains fidelity across both spatial and temporal levels of analysis, and thus may be valuable for examining relationships between networks and dynamic whole-brain architectures.

## Methods

### Subjects

Minimally preprocessed task-free fMRI data for 200 healthy adult humans (age 22-35, 112 female) were selected randomly from the Human Connectome Project S500 public data release^52^. Acquisition and analysis of data received institutional review board approval. Informed consent was obtained in accordance with the policies of the host institution, and data were de-identified prior to analysis.

### Image acquisition and preprocessing

Data were acquired on the 3T Connectom scanner (Siemens Healthcare, Erlangen, Germany) using multiband pulse sequences (*T*_*R*_ = 720 ms; *T*_*E*_ = 33.1 ms; 2.0 mm isotropic spatial resolution; multiband factor = 8)^53–56^. During task-free data acquisition, subjects were instructed to visually fixate on a crosshair. The data acquisition strategy is detailed elsewhere^57^.

Data were obtained as outputs of the Human Connectome Project’s denoising pipelines, detailed elsewhere^58^. In addition to standard fMRI preprocessing using FSL and FreeSurfer^59,60^ data were denoised to minimize the impact of subject movement on connectivity estimates. In brief, subject data were decomposed using ICA, and nuisance signals were removed via regression of realignment parameters, their temporal derivatives, and independent components identified as artifactual by a trained classifier (ICA-FIX^61,62^).

In order to divide the brain into functional parcels, we used group-level independent component analysis (1) to decompose the preprocessed images into 100 constituent signal sources common to all 200 subjects and (2) to identify where in the brain each signal was localized^34^. We then applied back-reconstruction (GICA1) to obtain, for each subject, 100 spatial maps and timeseries representing subject-specific analogues of each group-level independent component^63,64^. Component validity was assessed both qualitatively (via visual inspection) and via cross-correlation of component maps with canonical network maps^38^; components corresponding to movement or physiological noise were discarded, and 80 of the original 100 components were retained as functional parcels, or network nodes. The activation timeseries of each node was subject to additional preprocessing steps in GIFT, including demeaning and detrending, interpolation over artifact-related outliers (‘despiking’), removal of frequencies less than 0.01 Hz or greater than 0.15 Hz using a bandpass filter, and variance normalization of signal intensities.

### Canonical network discovery

The time-averaged functional connectivity between each pair of processed node timeseries was computed as the Pearson correlation coefficient^65^. This analysis yielded a symmetric, undirected graph with 3160 edges. The weight of the edge connecting node *n*_*i*_ to node *n*_*j*_ was encoded as feature *E*_*ij*_ in a symmetric 80 x 80 adjacency matrix *E*. This time-averaged connectivity matrix was used to separate nodes into canonical networks and to establish a reference against which transient connectivity metrics could be compared.

We applied a generalized Louvain-like community detection algorithm^36,37^ to a 70 x 70 submatrix of this adjacency matrix; this submatrix corresponded to cortical nodes and their connections. The Louvain resolution parameter was trained by performing community detection at a number of resolutions and penalizing the distance between the resultant partition and an established *a priori* partition^3^ (Supplementary Figure 1). Using this approach, we partitioned cortical nodes into six canonical networks.

### Dynamic functional connectivity and NC-state resolution

Dynamic functional connectivity among the 80 nodes was computed over a 44.64s tapered (rectangle convolved with a Gaussian) sliding window incremented 0.72s over 15-minute node timeseries^66^. In the absence of information about the timescale of dynamic fluctuations in connectivity, the probability of detecting such fluctuations in task-free data is optimized for a sliding window approximately 50s in duration^45^. The pairwise connectivity matrix during each time window was computed as a regularized precision matrix^28,67–69^. An aggressively denoised subsample of all data was selected by computing the mean framewise displacement^44^ during each time window and excluding any windows with a mean FD >0.18 mm; NC-state identification (as described below) was performed on the full sample and this subsample with comparable results.

We generated six network-specific graphs for each time window by extracting from the whole-brain graph only edges between nodes in the same network. Following an approach previously used to detect connectivity states^28^, we used k-means clustering (L1 distance) of these window-wise graphs to identify time windows during which each network exhibited relatively consistent connectivity patterns. We determined the number of clusters (connectivity patterns) for each network using a semi-formalized elbow criterion (Supplementary Figure 2). To ensure the validity of clustering, we performed clustering on data generated by independently permuting observations of each variable in order to preserve variable distributions without maintaining any explicit relationship between the variables. We found that the sum-of-squares error for the null data significantly exceeded that for the observed data (*p* < 0.01), a positive indication that a clustering approach was valid (Supplementary Figure 2C). We thus obtained for each network a set of cluster centroids along with a map assigning each time window to a centroid. We defined each centroid as an NC-state, or connectivity state. To ensure that NC-states represented dynamic reconfiguration within subjects rather than individual differences across subjects, the number of NC-states represented in each subject was computed. We also repeated connectivity state detection, as above, using all nodes in the cerebral cortex rather than only those assigned to a particular network.

To validate results, the clustering procedure was repeated on randomly selected split-half samples of the data. Similarity of subsample centroids was then assessed using the correlation distance metric (1−*r*, where *r* is the pairwise Pearson correlation between the connections of subsample centroids) as a proxy for similarity. In addition to the empirical split-half subsamples, clustering was performed on surrogate data (permuted split-half samples) generated by applying a random phase shift to the timeseries representing the strength of each connection within the network of interest over time. The similarity between empirical split-half centroids and analogous phase-shifted split-half centroids was then assessed using the correlation distance metric. Each centroid was reported as replicated if clustering of the empirical split-half samples produced centroids more similar to one another than to the centroids yielded by clustering the phase-shifted data (Supplementary Figure 3A). Overall, 36 of 46 centroids replicated, including all SAL and EXE NC-states, and all but two DAT and DMN NC-states. Qualitative visual inspection of all subsample centroids suggested replication rates similar to but greater than this automated approach.

### Contextual connectivity and concordance

For each subject, we used Louvain-like multilayer community detection to compute each node’s community membership at every point in time^37^. For each NC-state, we identified an average whole-brain connectivity context (WBCC) by computing a community-allegiance matrix over all time points in which the network exhibited the NC-state in question. In the allegiance matrix, an edge *E*_*ij*_ connecting nodes *n*_*i*_ and *n*_*j*_ is assigned a weight equal to the probability that *n*_*i*_ and *n*_*j*_ are assigned to the same community over the sampled time points (the “allegiance” of those nodes to one another)^70^. These context-specific allegiances were next quantified as a ‘displacement from baseline allegiance’, defined as the ratio of within- and between-network allegiances in a specific WBCC to the same within- and between-network allegiances averaged over all time. To facilitate visualization, these ratios were rescaled to represent relative shares of total allegiance in each WBCC and plotted on an exponential scale.

Additionally, a Bayesian ‘concordance’ metric was computed, which indexed the change in the probability that a particular network (or the whole cortex) exhibited a particular NC-state (or connectivity state) given information about the NC-state (or connectivity state) of another network (or the whole cortex):

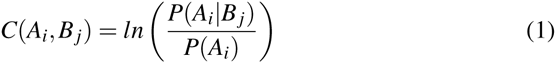

wherein for each pair of network-specific NC-states, *A*_i_ represents NC-state *i* of network A; *B*_*j*_ represents NC-state *j* of network B, where *j* may equal *i*

Concordance was zero-centred by applying a logarithm to the posterior-to-prior probability ratio; positive concordances thus corresponded to states more likely to co-occur than predicted by chance, while negative concordances corresponded to states less likely to co-occur than predicted by chance.

Three null models were used to generate control distributions of concordance data under the assumption of complete independence of the networks. These were generated by shuffling or simulating the assignment of each time point to a particular set of NC-states. In the first model, the observed NC-state assignments for each network were randomly permuted across subjects. In this way, the observed trends of occurrence of connectivity states over time was preserved, but any explicit relationship between NC-states in different networks was abolished. In the second model, NC-state assignments were simulated using the observed initial conditions and Markov chain transition models computed from the observed data. In the third model, the phase of the observed NC-state assignments was randomly shifted. In this way, any apparent concordance that was attributable to static individual differences (or to sampling variability) was preserved, but dynamic concordances were abolished. A concordance matrix was then generated, as above, for the permuted or simulated data. Null distributions for hypothesis testing were generated from 100 repetitions of each null model. If a concordance was non-significant under any of the three null models, then it was marked as non-significant.

### Subsystem identification

Four NC-states corresponding to fractionation of a brain network into subsystems were identified through a qualitative screen. Dynamically engaged subsystems of brain networks were then identified through hierarchical clustering of the average connectivity profiles of all nodes in these NC-states. Hierarchical clustering was performed using the correlation distance metric, defined as 1−*r*, where *r* is the Pearson correlation coefficient between the compared connectivity profiles. Subsystems of brain networks were considered to be recruited in a particular NC-state if the correlation distance between the connectivity profiles of their constituent nodes did not exceed 0.4.

### Evaluation of null/independence hypothesis

We evaluated the null hypothesis of complete network independence by comparing the empirical contexts of each NC-state to phase-randomized contexts of each NC-state. For each network, we applied a randomized phase shift to the timeseries representing the strength of each connection over time^28,45^ (the edge-weight timeseries). Only connections outside of the network of interest were phase-shifted in this manner; thus, dynamic structure was preserved within each network but abolished for the remainder of the brain. Contexts were then computed for each NC-state for the phase-randomized data, as described above.

Our analysis was predicated upon the following assumptions:

- Randomly phase-shifting each edge-weight timeseries preserves the mean and variance of the timeseries, but disrupts the overall dynamic covariance structure^28^, which is dependent upon common features across multiple edge-weight timeseries.
- If a network’s intrinsic connectivity state were independent of its whole-brain environment, then the network’s states would not co-occur with any consistent changes in the covariance structure of the remainder of the brain.
- If networks are completely independent, then a randomized phase shift of all edge-weight timeseries not in the network of interest will not meaningfully change the similarity between contexts.

When we applied a randomized phase-shift to all edge weight timeseries outside of a network of interest, we instead observed that the resultant phase-shifted NC-state contexts were, without exception, more similar to one another than were the empirical contexts of NC-states (*p* << 0.001, paired Wilcoxon signed-rank test), suggesting that NC-states occurred in the context of specific changes in the whole-brain connectivity structure (Supplementary Figure 3B).

### Independence

The independence of each canonical network was computed using two metrics: mutual variation and mean concordance. For each network, null distributions for each metric were generated on the basis of 50 *pseudo-networks*. Each pseudo-network was defined to include the same number of nodes and NC-states as the canonical network in question. However, unlike the case for canonical networks, nodes were assigned to pseudo-networks randomly and not on the basis of community structure or previous scientific results. Pseudo-network NC-states were then computed using k-means clustering in a manner analogous to canonical network NC-states. For each canonical and pseudo-network NC-state, contexts were obtained (1) as allegiance matrices, defined as the probability that each pair of nodes would be assigned to same community while that particular NC-state was present, and (2) as contextual centroids, defined as the mean of all Fisher-transformed whole-cortex windows during which a network or pseudo-network expressed that NC-state.

For independence analysis using the mutual variation metric, contextual centroids were treated as a proposed clustering solution for the entire cortex, with the understanding that a network that was less independent of the whole cortex would provide a better clustering solution for the cortex. The goodness of this clustering solution was thus computed as a proxy for the network’s independence from the cortex as follows. The correlation distance from each z-transformed whole-cortex window to the nearest contextual centroid was computed (the “within-cluster” distance). All distances were squared and subsequently added together to determine a within-cluster sum-of-squares (WCSS) error term. The WCSS reflected the extent to which the proposed clustering solution (which was based only on information about the temporal variance of a single canonical or pseudo-network) explained the temporal variance present in the entire cortex; a lower error term corresponded to a better clustering solution and thus to less independence.

The theoretical upper limit on clustering efficacy (maximal interdependence) corresponded to the clustering solution that minimized the WCSS error; this was obtained by clustering all cortical features (taking into account the temporal variance of the entire cortex rather than only that of its subsystems). The theoretical lower limit on cluster efficacy (maximal independence) would result in maximization of the WCSS error and corresponded to centroids identical to the time-averaged cortical connectivity except in the network of interest, where they were identical to the network’s NC-state centroids. Tests for significance (two-sided Wilcoxon signed-rank tests) were performed for each canonical network’s WCSS error score relative to the null distribution generated from the 50 pseudo-networks with similar properties. Following the independence analysis, an independence score was obtained for each network by scaling the mean WCSS error among pseudo-networks to zero, maximal interdependence to -1, and maximal independence to 1.

A second assay for independence was performed using the concordance metric. Here, interdependence was operationalized as the concordance of (pseudo-)network NC-states with 8 whole-cortex states, with positive and negative concordances computed separately because of their different interpretations. More interdependent systems were predicted to exhibit greater positive and negative concordances. The mean positive and negative concordances between (pseudo-)network NC-states and whole-cortex states were separately computed. Results using this metric (with the exception of those for the somatomotor network) were convergent with the results from the WCSS approach.

### Identification of a highly interdependent system

The “interdependent” (INT) system was identified through a two-step process. First, a tally was obtained of the frequency with which nodes appeared in pseudo-networks that scored in the bottom quintile of WCSS errors. A subset of nodes that frequently often occurred in such “interdependent” networks was thus identified, and pseudo-network generation (8 nodes, 8 NC-states) was repeated 100 times, with random drawing from only this subset of nodes. On the whole, the resultant pseudo-networks exhibited notably lower WCSS errors than did those selected from all nodes. Among these pseudo-networks, the one with greatest interdependence (lowest WCSS error) was selected as the INT system. Interdependence was re-evaluated and reproduced in split-half samples and with 16 NC-states and cortical states instead of 8. Independence was also evaluated for pseudo-networks generated from nodes with (1) maximal participation coefficient and (2) mean allegiance to nodes other than canonical partners without significant results.

## Acknowledgements

R.C. and A.B.S. acknowledge funding support from the Pomona College Department of Neuroscience. Data were provided by the Human Connectome Project, WU-Minn Consortium (Principal investigators: David Van Essen and Kamil Ugurbil; 1U54MH091657) funded by the 16 NIH institutes and centres that support the NIH Blueprint for Neuroscience Research; and by the McDonnell Center for Systems Neuroscience at Washington University.

## Author contributions

R.C. and A.B.S. formulated the project; R.C. designed the analyses; R.C. and J.S.N. performed the analyses; R.C. and A.B.S. drafted the manuscript with revisions from J.S.N. and L.Q.U.

## Additional information

The authors have no competing financial interests to declare.

## Supplementary Information

### Supplementary Figures

**Supplementary Figure 1.**
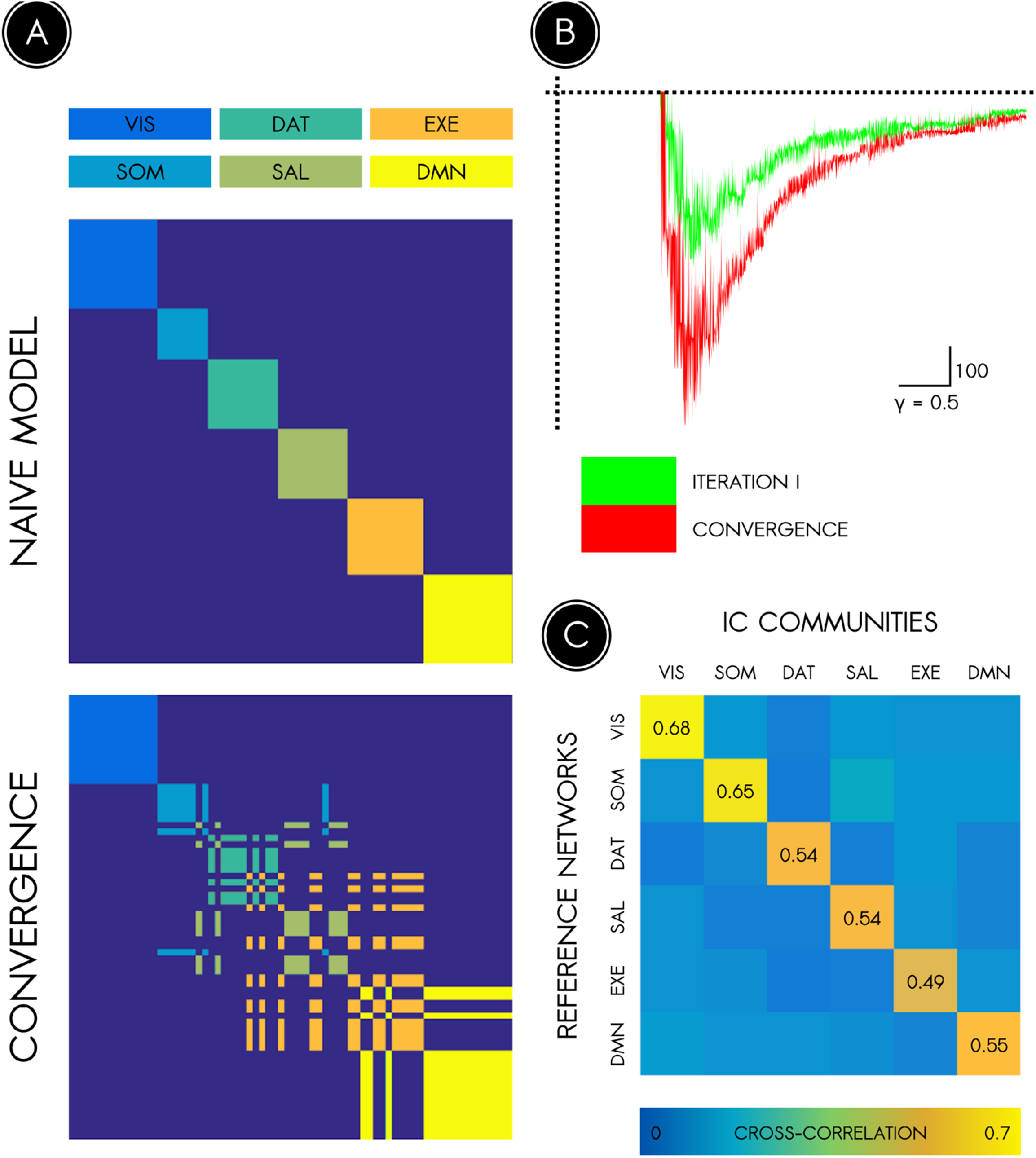
Training the resolution parameter of the generalized Louvain algorithm to discover canonical networks. An existing 7-network partition of the cerebral cortex (Yeo et al., 2011) was used to train the resolution parameter of the generalized Louvain algorithm. **(A)** *Top*, the spatial model derived from cross-correlations of each community with the *a priori* partition, which served as a starting point for training the algorithm. *Bottom*, when the algorithm converged, the Louvain solution and spatial model were identical. **(B)** The distance between the Louvain partition and the spatial model across a range of values of the resolution parameter gamma. The optimal solution consistently occurred at a value of gamma near 1.3. The observed instability along the ordinate occurs due to degeneracy of the Louvain partition. **(C)** Spatial cross-correlations of communities of nodes with reference networks from the *a priori* partition ranged from approximately 0.5 to 0.7, establishing a one-to-one correspondence between our communities and the canonical reference networks.

**Supplementary Figure 2.**
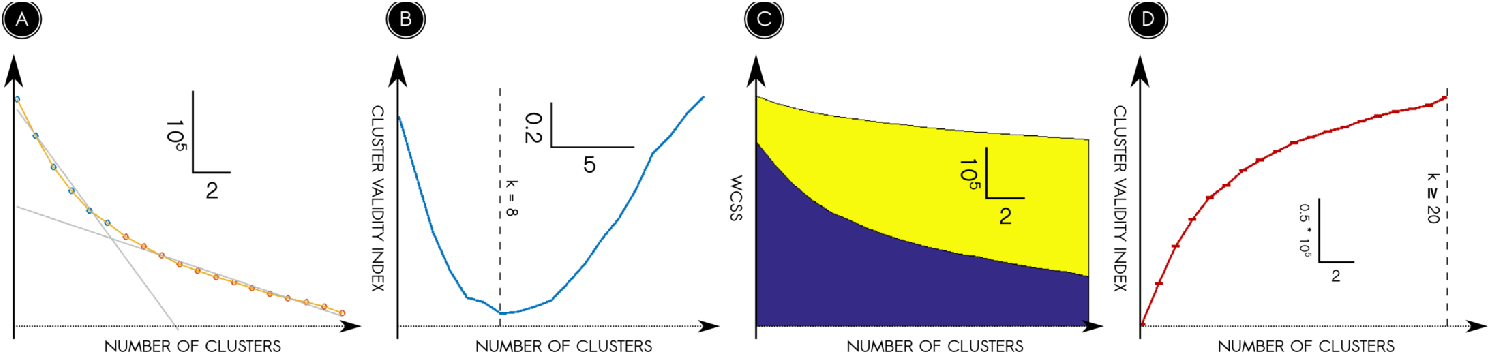
Validity of clustering-based data reduction is established via permutation. **(A)** Sample clustering validity plot for the salience network, depicting the elbow computation for the optimal solution. Two least-squares lines were computed, with the putative solution demarcating the point of separation along the abscissa between data included in each computation. The putative solution that optimised fit for both lines was selected. **(B)** A semi-formalized elbow criterion suggested an optimal solution of *k* = 8 clusters. **(C)** The empirical clustering validity plot for the salience network (blue) in comparison with the mean null clustering validity plot (yellow) across a range of *k*. **(D)** A formalized gap criterion suggested an optimal solution of at least 20 clusters. Error bars indicate standard deviation.

**Supplementary Figure 3.**
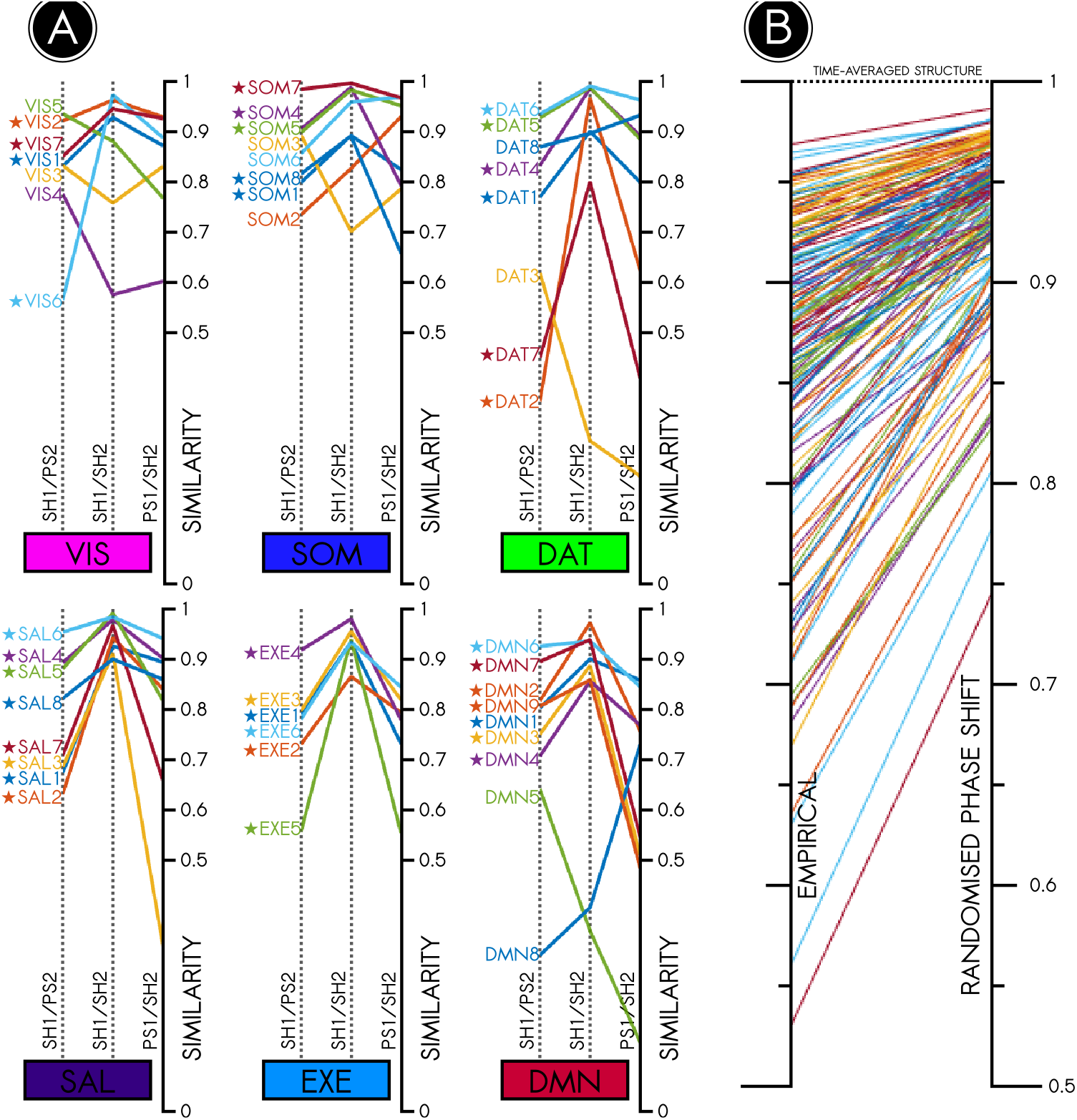
Randomized phase shifts of edge-weight timeseries establish NC-state replication and network interdependence. **(A)** To determine whether each NC-state replicated across randomized split-half subsamples, we compared the similarity of empirical split-half NC-states to one another (SH1/SH2) against the similarity of empirical split-half NC-states to phase-randomized split-half NC-states (SH1/PS2 and PS1/SH2). If similarity between empirical NC-states was greater than similarity of empirical NC-states to phase-randomized NC-states, this was a positive indication that the NC-state replicated. NC-states denoted with a star (★) are those that replicated according to this criterion. In total, 36 out of 46 total NC-states replicated. Among these were all SAL and EXE NC-states, all but 2 NC-states of DAT and DMN each, and the majority of VIS and SOM NC-states. The overall featurewise similarity between split-half NC-states was approximately 0.88. **(B)** We formally evaluated whether a network’s intrinsic state was independent of its context by comparing (i) the similarity of empirically observed contexts to one another against (ii) the similarity of phase-randomized contexts to one another. (If all contexts recapitulated the time-averaged connectivity, then their similarity coefficients would approach a maximal value of 1.) Phase-randomizing extrinsic connections resulted without exception in contexts that were more similar to one another than the empirically observed contexts, indicating that NC-states empirically occurred in specific whole-brain contexts (*p* << 0.001, paired Wilcoxon signed-rank test).

**Supplementary Figure 4.**
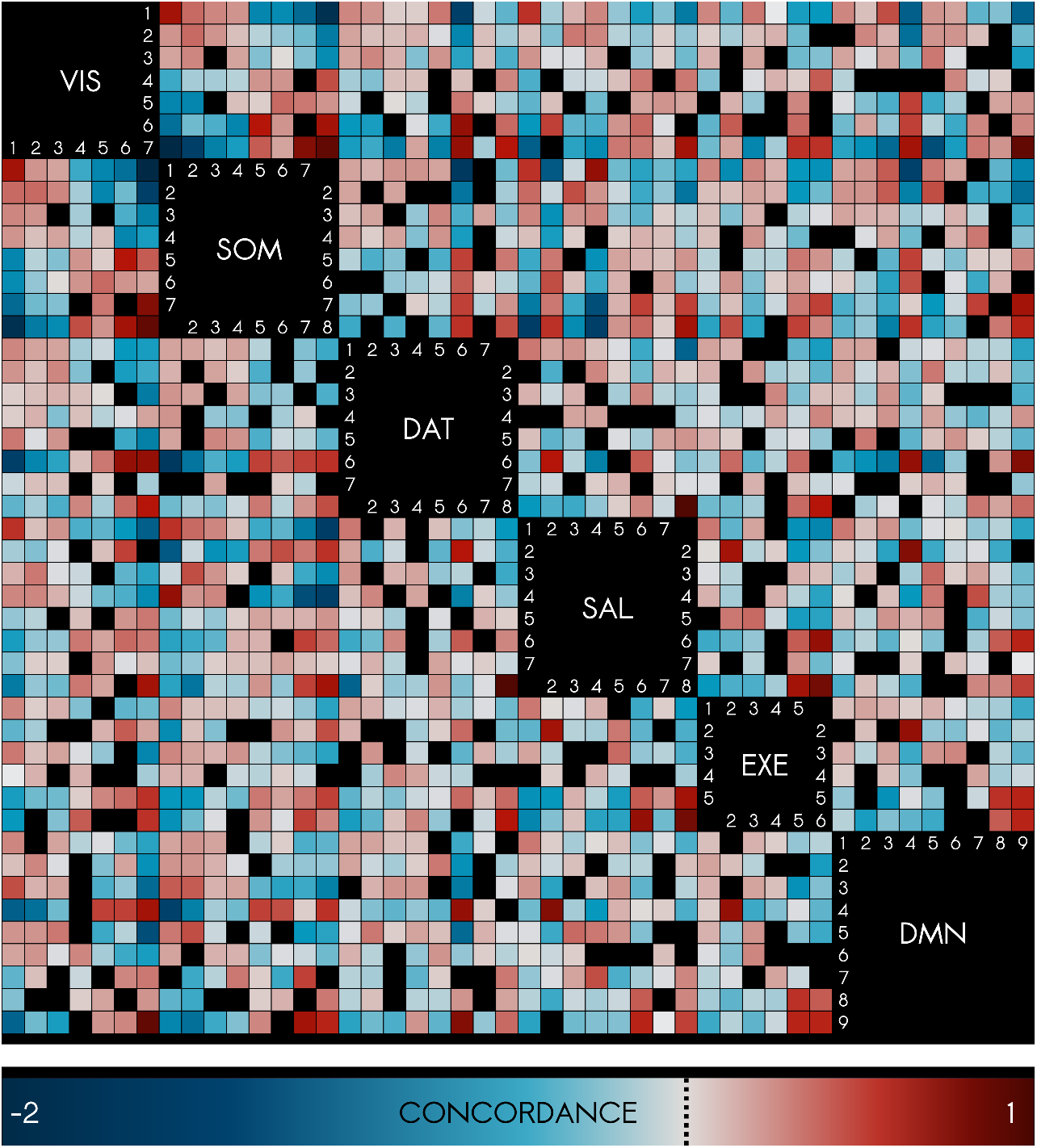
Concordance matrix across NC-states: Functional interdependence of networks.. The figure illustrates pervasive interdependence between canonical networks when examined at the level of NC-states. Cooperativity between brain systems is revealed in a state-by-state matrix of Bayesian concordances, which captures co-occurrences of single-system connectivity patterns that deviate from the prior probability derived under complete independence of brain systems. Compared with permuted and simulated null models assuming independence, nearly all NC-states exhibited a significant degree of concordance or discordance (*p* < 0.01, two-sided Wilcoxon signed rank test, Bonferroni corrected). The fewer non-significant concordances are blacked out for ease of visualization.

**Supplementary Figure 5.**
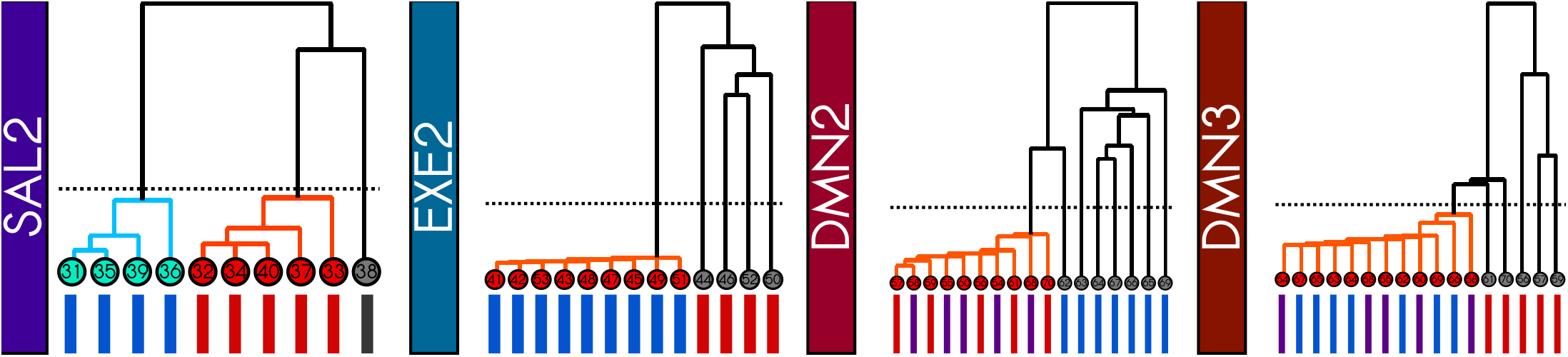
Criterion for subnetwork identification. To identify dynamic subnetworks, hierarchical clustering of whole-brain node allegiance profiles was performed. The resultant dendrograms are shown here for the states displayed in Figure 6. A hard cut-off at a correlation distance of 0.4 was used to define cohesive subnetworks. While not cohesive, nodes of the auxiliary executive subsystem did display some common changes in overall connectivity.

